# A scalable, multi-resolution consensus clustering approach for prioritising robust signals from high-throughput screens

**DOI:** 10.1101/2024.09.05.611300

**Authors:** Shaine Chenxin Bao, Kathleen I. Pishas, Karla J Cowley, Qiongyi Zhao, Emily C. A. Goodall, Ian R. Henderson, Evanny Marinovic, Mark S. Carey, Ian G. Campbell, Kaylene J. Simpson, Dane Cheasley, Dalia Mizikovsky, Nathan J. Palpant

**Affiliations:** Institute for Molecular Bioscience, The University of Queensland, Brisbane, Australia; Peter MacCallum Cancer Centre, Melbourne Australia; Victorian Centre for Functional Genomics, Peter MacCallum Cancer Centre, Melbourne, Australia; Division of Gynecology Oncology, Department of Obstetrics and Gynecology, University of British Columbia, Vancouver, British Columbia, V5Z 1M9, Canada; Department of Clinical Research, BC Cancer, Vancouver, British Columbia, V5Z 4E6, Canada; The Sir Peter MacCallum Department of Oncology, The University of Melbourne, Parkville, Australia; Department of Biochemistry and Pharmacology, University of Melbourne, Parkville, Australia

## Abstract

Modern biology increasingly relies on large-scale screening to generate high dimensional datasets with potential to accelerate discovery. However, analysing these complex datasets remains challenging, particularly in applications where the underlying structure and groupings are unknown, and high dimensionality introduces noise and artifacts that make follow up studies difficult to prioritise. Here, we present an unsupervised consensus clustering tool that quantifies biologically meaningful patterns based on multi-scale data organisation to guide decision-making in high-throughput screening. Using large-scale drug screening data in cancer cell lines and bacterium model, we demonstrate its ability to use diverse data inputs to prioritize robust drug clusters with shared biological mechanisms and conserved drug responses. This method addresses key limitations associated with prioritising robust, actionable hits from scalable screening data.

---

Advancements in high-throughput sequencing technologies have generated unprecedented volumes of biological data^1,2^. Traditional comparative analyses that require groups to be labelled *a priori* are inadequate for such complex datasets. Unsupervised clustering methods are therefore ideal to assist in identifying hidden structure in high-dimensional feature spaces without prior assumptions. Clustering samples based on their molecular or phenotypic features enables data-driven approaches to discover meaningful relationships or shared mechanisms. Despite this, clustering high dimensional data remains challenging, particularly in applications where the number of true biological groupings are unknown and the signal is confounded by stochastic noise^3^.

Consensus clustering methods have emerged as a solution to improve clustering sensitivity and robustness, particularly in the single cell and bulk transcriptomics field^4,5^. These approaches assume that true biological clusters remain stable despite varied hyperparameters or algorithms^6^. Rather than relying on a single clustering result, they aggregate multiple clustering solutions to build a similarity matrix representing how often objects co-cluster across iterations and then perform clustering on the compiled consensus matrix^7^. Despite improving overall cluster accuracy, existing methods do not provide a systematic approach to evaluate cluster quality^5,8-13^. Without an efficient way to determine the true underlying biological patterns and prioritise actionable data clusters, follow-up analysis becomes resource intensive and risky. Furthermore, clustering performance is inherently limited by the data quality, which can suffer in large-scale screens, making it essential to detect low-quality, stochastic clusters. In this study, we present UnTANGLeD, an unsupervised consensus clustering pipeline that identifies robust biological patterns in high-dimensional data with post-hoc evaluation to quantifiably prioritise clusters for actionable and efficient follow-up.

UnTANGLeD takes any dimensionality-reduced dataset following pre-processing and performs iterative clustering to evaluate signatures of similarity across increasing granularity. The co-clustering frequencies of object pairs are used to construct a consensus matrix, transforming raw distance into an interpretable robustness-based metric quantifying relationship consistency. By leveraging multi-resolution clustering, this method effectively denoises data, retaining genuine biological relationships that persist across multiple data scales while filtering out weak or spurious associations. It further employs a stability-driven approach to determine the optimal number of clusters that capture maximal information from the data. After hierarchical clustering, clusters are stratified based on internal coherence (correlation), robustness (silhouette score), and assessed for conservation across biological contexts. This provides researchers with an unbiased framework for prioritizing robust and biologically meaningful clusters for further investigation, optimizing resource allocation in screening scenarios with limited or no prior annotation (**Figure 1a**). Originally developed to identify gene programs from sparse, gene-trait association data^14^, we have established UnTANGLeD as a versatile pipeline for post-screening analysis integrating clustering, cluster optimisation, evaluation, and conservation metrics in one workflow applicable to diverse data modalities and experimental designs. In this study, we aim to demonstrate its utility across various large-scale screens, including a high-content imaging drug screen of ovarian cancer cell lines^15^, a gene deletion library in *E. coli*^16^, and profiling of transcriptional responses to drug perturbations^1^(**Figure 1b**).

**Figure 1:**
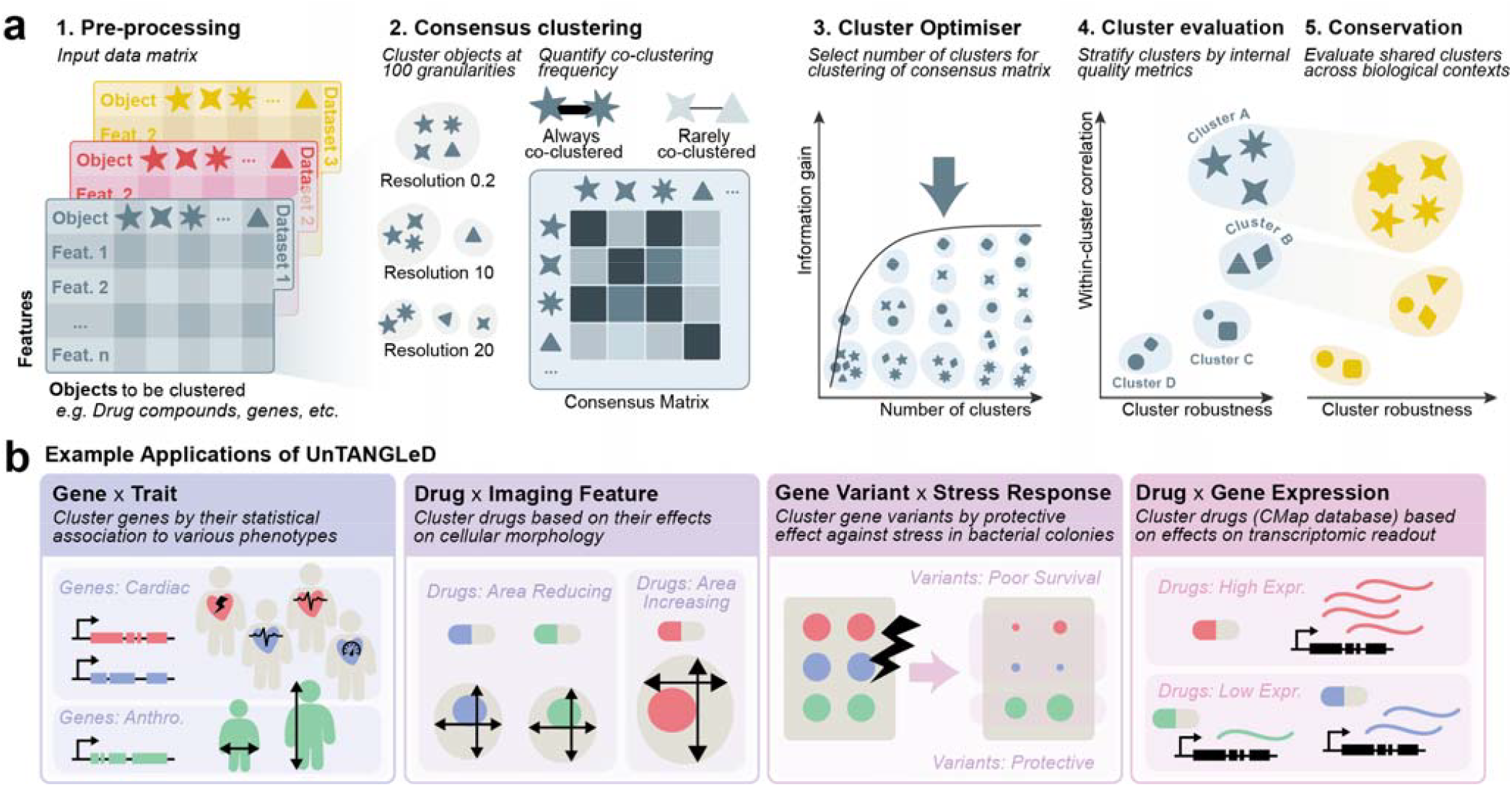
Unsupervised clustering prioritisation pipeline and application of UnTANGLeD across different data modalities. **(a)** Schematic overview of the UnTANGLeD workflow. **(b)** Representative applications of UnTANGLeD across diverse biological data modalities.

First, we analysed high content morphology imaging data measuring the phenotypic response of three low grade serous ovarian carcinoma cell lines with unique genetic profiles (AOCS-2, VOA-6406, SLC58), and one normal ovarian surface epithelial cell line (IOSE-523) exposed to a diverse library of 5,596 drugs, including FDA-approved drugs, kinase inhibitors, methylation modulators, and investigational agents, at 3 different concentrations (10, 1, 0.1µM) (**Figure 2a**)^15^. We tested the utility for UnTANGLeD to leverage the 2,175 imaging features to cluster over 16,750 drug conditions by their shared morphological changes in cells and prioritise robust drug clusters with shared biological mechanisms. Raw datasets were pre-processed and reduced in dimensionality using principal component analysis. To assess clustering stability across different granularities, 100 iterations of Seurat’s shared nearest neighbour (SNN) algorithm were run at increasing resolutions. These results were combined into a consensus matrix, where each element represents the co-clustering frequency of any two drugs across all resolutions. This re-configures the high-dimensional drug-by-imaging matrix into a sparse consensus matrix that quantifies the relationship strength between all drug pairs (**Figure 2b**). UnTANGLeD then performs agglomerative clustering using Ward’s minimum variance on the consensus matrix from 2 to 300 clusters, calculating the average silhouette score for each number of clusters as a metric of cluster quality across the data set. The silhouette score (−1 to 1) assesses how similar each object is to other objects in its assigned cluster and how different it is from objects in the nearest neighbouring cluster. As the silhouette score is calculated on the consensus matrix, a high silhouette score indicates a stable, distinct grouping. When increasing the number of clusters no longer improves the average silhouette score, no more biologically distinctive groups can be gained by further subdividing the data. In our analysis, the optimal number of clusters was defined at approximately 200 clusters across all four cell lines (**Figure 2c**).

**Figure 2:**
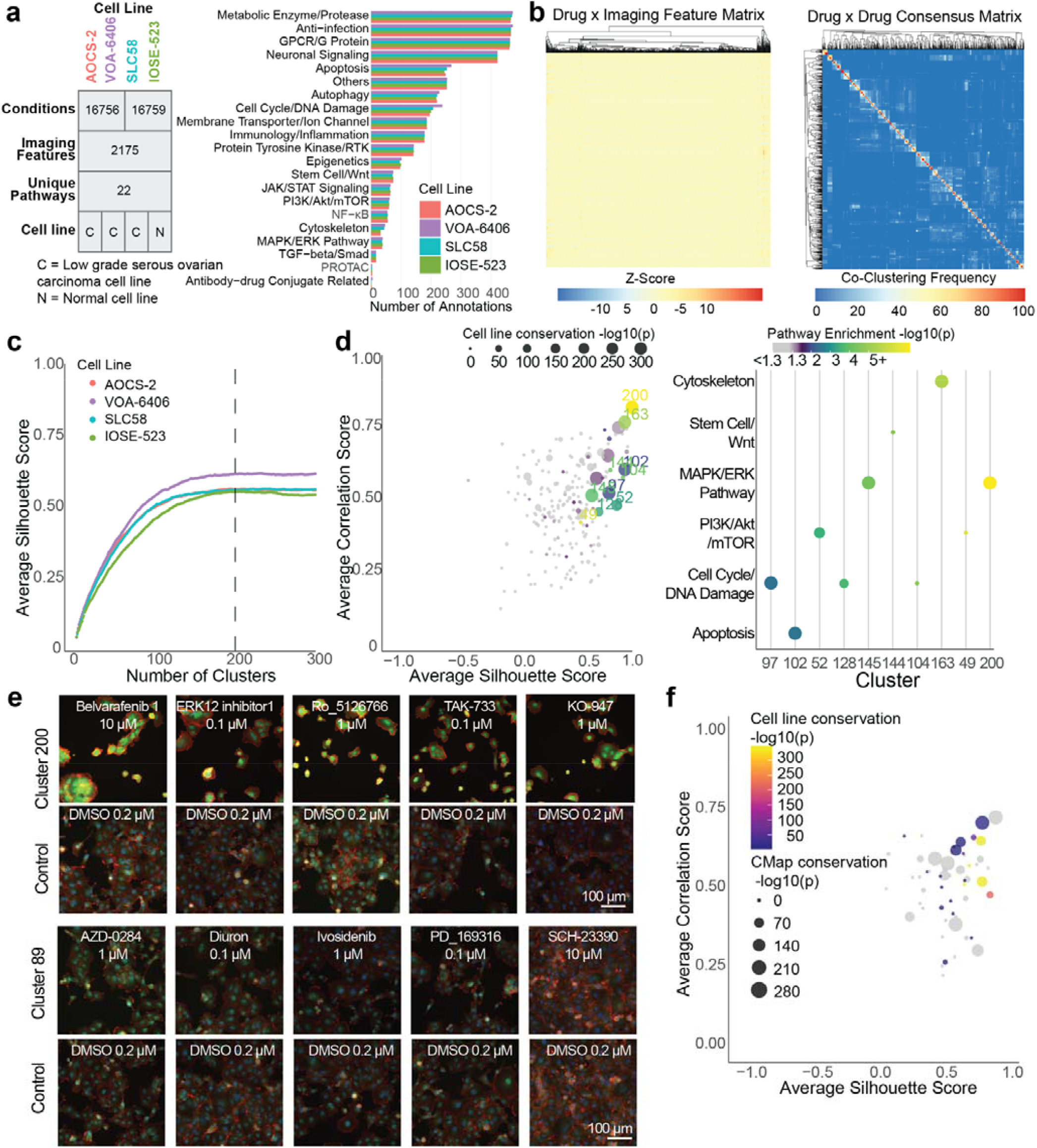
Unsupervised clustering reveals biologically meaningful drug clusters based on cellular morphology. (**a**) Dataset characteristics for the four cell lines analysed (3 genetically variable ovarian cancer: AOCS-2, VOA-6406, SLC58; 1 normal: IOSE-523). Each cell line was treated with a 5596-compound library for 72 hours. The table shows the number of unique conditions (compound at varying concentrations), imaging features initially extracted, unique pathways. Bar plot displays the distribution of annotated biological pathways across all compounds. (**b**) Transformation of high-dimensional data into a sparse similarity matrix. Heatmaps of a subset of the dense, raw drug-by-imaging matrix (top) and the sparse drug-by-drug consensus matrix (bottom). The consensus matrix represents drug co-clustering frequency across 100 iterations of Seurat clustering at resolutions 0.2 to 20 with an interval of 0.2. (**c**) Determination of optimal cluster number using average silhouette scores across 2-300 clusters for all four cell lines. ~200 clusters were selected for all cell lines based on the average silhouette score plateau point (indicated by dashed line). (**d)** Scatter plot of clusters in cell line SLC58 stratified by average cluster silhouette score and correlation score. Pathway annotations for the top 10 clusters with the highest pathway enrichment −log10 p-adjust for cell line SLC58. Size indicates conservation significance (Conservation −log10 p-adjust) across cell lines while colour represents pathway enrichment significance (Enrichment −log10 p-adjust). (**e**) Representative microscopy images of cells (1 field per well) from cell line SLC58 treated with selected drug conditions from example top-ranked Cluster 200 and bottom-ranked Cluster 89. Images shown with DMSO control and 100μm scale bar. Cells were stained with DAPI (nucleus; blue), CellMask (plasma membrane; green) and phalloidin/rhodamine (F-actin; red; red). **(f)** Orthogonal validation of clusters using transcriptomic data. Scatter plot of UnTANGLeD clusters stratified by average silhouette score and correlation score. Size indicates −log10 p.adjust conservation significance of UnTANGLeD drug clusters with those derived from Connectivity Map (CMap) using gene expression, limited to clusters containing more than 5 overlapping drugs.

Clusters were next stratified by the silhouette score, indicative of cluster robustness, and the average pairwise correlation of imaging features within each cluster, indicative of intra-cluster similarity (**Extended Figure 1a-d**). Given the genetic heterogeneity of ovarian cancer, we were interested in identifying drug mechanisms that induce consistent phenotypic responses robust to genetic variation. To assess this, we evaluated the consistency of drug clusters across cell lines using three conservation metrics. Many of the top ranked drug clusters, defined as those scoring high on average silhouette and correlation scores, were significantly conserved across all four ovarian cell lines (**Figure 2d**). To assess the biological significance of identified clusters, we performed hypergeometric tests to evaluate the enrichment of annotated pathways for each drug cluster. Across all four cell lines, clusters were significantly enriched for cancer-related pathways (**Extended Figure 1e-h**). For instance, the top ranked drug clusters from cell line SLC58, were strongly enriched for key oncogenic pathways including *MAPK/ERK*^*17*^, *PI3K/AKT/mTOR*^*18*^, and Cell cycle/DNA damage^19^ (**Figure 2d**). To reinforce these findings, we examined representative microscopy images of cells treated with drugs from selected clusters. Top-ranked clusters exhibited more consistent morphological changes in treated cells vs DMSO control, contrasting with inconsistent morphologies in poorly ranked clusters (**Extended Figure 2**). Notably, the observed morphological changes closely aligned with the enriched pathways for these clusters. For example, the drugs in top-ranked Cluster 200 from cell line SLC58 was enriched for MAPK/ERK pathway and showed morphological changes including cell shrinkage, membrane blebbing, and fragmented nuclei, consistent with apoptosis induction (**Figure 2e**). These observations align with the known role of *MAPK/ERK* pathway in regulating cell proliferation and survival, particularly in low-grade serous ovarian cancer pathogenesis^17,20^, where its disruption can lead to apoptosis^21^. We further validated our clustering outputs using Connectivity Map (CMap)^1^, an independent dataset characterising changes in gene expression in response to drug perturbations, focusing on breast, prostate cancer and leukaemia cell lines. This comparison allowed us to evaluate whether the morphological signatures identified by imaging align with transcriptomic changes induced by the same drugs. UnTANGLeD was applied to CMap to yield 175 drug clusters. Next, we assessed the conservation between these transcriptomics-based clusters and our imaging-based clusters across all four ovarian cell lines, focusing on drugs present in both datasets (**Extended Figure 3a**). Indeed, overlapping drug sets between the two modalities were conserved, with stronger conservation observed in highly ranked clusters (**Figure 2f, Extended Figure 3b)**, implying concordant changes at the transcriptomic level and providing orthogonal validation to our clustering.

To contextualize UnTANGLeD’s performance and its advantages, we benchmarked it against two widely used traditional clustering approaches, k-means and hierarchical clustering, on dimensionality reduced data. K-means iteratively assigns data points to their nearest cluster centres and recalculates centres until convergence, while hierarchical clustering progressively merges similar points or clusters based on a defined distance metric. While attempting to identify optimal cluster numbers using silhouette scores, we found that both methods’ scores rapidly fell and plateaued near 0, despite dimensionality reduction of the dataset, indicating their inability to maintain or quantify meaningful cluster separation (**Figure 3a, Extended Figure 4a**). By contrast, UnTANGLeD demonstrated the ability to improve clustering quality with increasing cluster granularity. For comparison, we set the cluster number to 200 for all methods based on the previously determined optimal cluster number and evaluated the resulting clusters based on internal quality metrics (silhouette and correlation scores), biological relevance (pathway enrichment), and shared drug mechanism (cell-line conservation). UnTANGLeD consistently produced a clear stratification pattern where clusters in the upper right quadrant with both high internal coherence and distinctiveness showed stronger biological relevance, evidenced by increased pathway enrichment and cross-cell line conservation (**Figure 3b, Extended Figure 1a-d**). K-means clusters showed uniformly poor silhouette scores despite varying correlation values, indicating an inability to form biologically distinct clusters (**Extended Figure 4b)**. Unlike UnTANGLeD and k-means which produced normally distributed cluster size (**Figure 3c**), hierarchical clustering produced either very small, highly correlated clusters due to feature redundancy or large, catch-all clusters with decreased robustness (**Extended Figure 4c, Extended Figure 5**). In addition, neither silhouette nor correlation scores effectively stratified clusters by biological relevance for both k-means and hierarchical clustering (**Extended Figure 6**).

**Figure 3:**
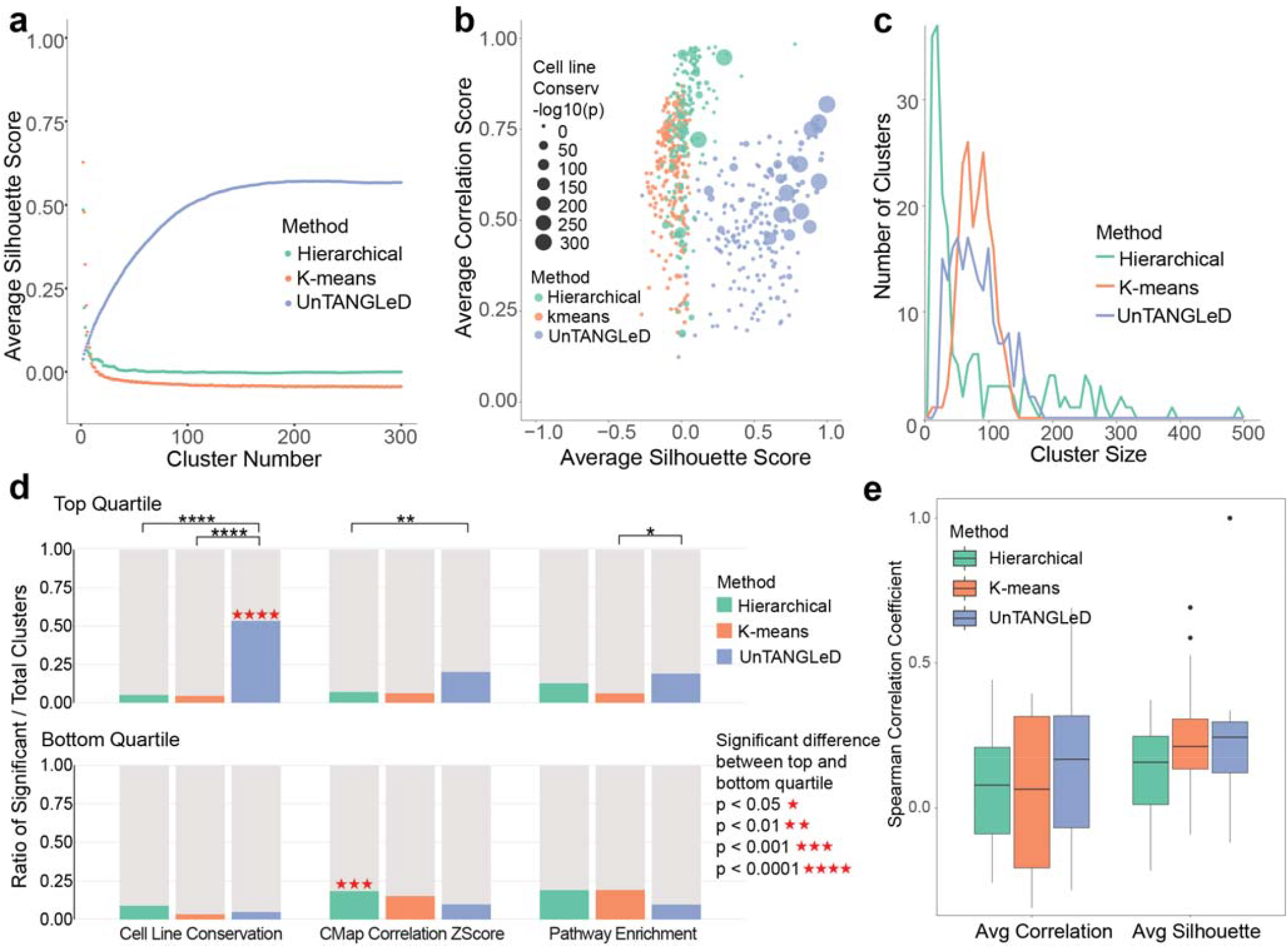
UnTANGLeD outperforms traditional clustering methods in prioritising biologically robust drug clusters. **(a)** Comparison of average silhouette scores across increasing cluster numbers (2-300) for UnTANGLeD (blue), k-means (orange), and hierarchical (green) clustering. **(b)** Scatter plot comparing cluster quality metrics (average silhouette score vs. average correlation score) for UnTANGLeD (blue), K-means (orange), and hierarchical (green) clustering. Dot size indicates conservation significance across cell lines (-log10 p-adjust). **(c)** Distribution of cluster sizes for UnTANGLeD (blue), k-means (orange), and hierarchical clustering (green). **(b-c)** Data shown for SLC58 cell line. **(d)** Comparison of significant cluster ratios compiled from four cell lines combined across UnTANGLeD (blue), k-means (orange), and hierarchical (green) clustering for three biological relevance metrics. Top quartile shows top-performing clusters (>75% quantile) while bottom quartile shows bottom-performing clusters (<25% quantile), as defined by 75^th^ of 25^th^ percentile of average correlation and silhouette scores for each individual cell line. Biological significance is defined as followed: CMap correlation Z-score (>2), conservation (adj. p-value <0.05), and pathway enrichment (adj. p-value <0.05). Statistical significance between methods and between top and bottom quartiles was calculated using Fisher’s exact test (*p<0.05, **p<0.01, ***p<0.001, ****p<0.0001). **(e)** Boxplots comparing the distribution of Spearman correlation coefficients between internal quality metrics (silhouette score, correlation score) with biological relevance metrics (pathway enrichment and cell line conservation) for UnTANGLeD (blue), K-means (orange) and Hierarchical clustering (green). Plotted with median, interquartile range and outliers.

To assess each method’s ability to prioritize biologically meaningful clusters, we first examined the average significance for clusters showing significant pathway enrichment and cell-line conservation. As shown in **Extended Figure 4d-e**, UnTANGLeD clusters on average demonstrated greater conservation across genetically diverse ovarian cancer cell lines compared to k-means and hierarchical clustering, though statistical significance was reached in cell line VOA-6406 only. While aggregated performance metrics across all clusters showed modest differences between methods, we hypothesized that biological relevance would not be uniformly distributed across all clusters but would correlate with internal quality metrics. To test this, we stratified clusters into top (highest 25%) and bottom (lowest 25%) quartiles based on their average silhouette and correlation scores. We then performed Fisher’s exact tests to assess whether the proportions of biologically significant clusters differed between clustering methods (**Extended Table 1a-b)**. This analysis revealed that UnTANGLeD’s top-ranked clusters contained significantly more cell-line conserved clusters compared to those generated by k-means and hierarchical clustering (OR <= 0.1, p < 1.3×10^−10^) (**Figure 3d**), highlighting its ability to identify consistent drug responses despite genetic heterogeneity between cell lines. Significantly more UnTANGLeD top-ranked clusters showed stronger transcriptional correlation in the CMap dataset compared to hierarchical clustering (OR = 0.05, p = 3×10□□), with a modest advantage over k-means (OR = 0.71, p = 0.4), indicating morphology-based drug clusters prioritised by UnTANGLeD tend to induce similar gene expression patterns. UnTANGLeD had significantly more pathway enriched clusters in its top-ranked clusters compared to k-means (OR = 0.28, p = 0.038) and a similar proportion compared to hierarchical clustering (OR = 0.99, p = 1.0) (**Figure 3d**). When comparing between upper and bottom quartile (**Extended Table 2**), UnTANGLeD had significantly more cell-line conserved clusters in upper quartile while no significance is observed for k-means and hierarchical clustering (**Figure 3d**). Most importantly, UnTANGLeD’s internal quality metrics correlation (Spearman’s ρ = 0.168) and silhouette (Spearman’s ρ = 0.277) scores aligned more strongly with external biological validation measures (pathway enrichment and cross-cell line conservation) on a continuous scale **(Figure 3e)**. This enables more confident prioritization of promising drug clusters for downstream investigation without requiring extensive *a priori* biological knowledge. Collectively, these data demonstrate that UnTANGLeD effectively parses large-scale morphology data to reveal robust, meaningful drug groupings characterized by strong pathway enrichment, high cross-cell line conservation, and alignment with orthogonal gene expression data. By grouping drugs by their phenotypic effects on fundamental cancer mechanisms across genetically distinct ovarian cell lines, UnTANGLeD prioritises robust drug clusters with metrics that help inform downstream decision making.

To further illustrate its utility, we evaluated UnTANGLeD on two additional high dimension screening data. First, applied to a genome-wide *E. coli* single gene-deletion library screened against 324 stress conditions^16^, UnTANGLeD effectively grouped gene-deletion mutants based on shared phenotypic responses, stratifying highly robust gene clusters enriched for shared biological processes to environmental stresses. It similarly identified condition clusters with similar gene signatures across mutants, prioritizing clusters with increased drug target enrichment (**Extended Figure 7**). Likewise, in analysing CMap’s transcriptional responses to over 1,000 drug perturbations across three other tumour cell lines (Luminal A breast cancer, prostate cancer and leukaemia)^1^, UnTANGLeD identified highly conserved drug clusters with shared mechanisms of action and drug indication enrichment based on their perturbed gene expression signatures (**Extended Figure 8**).

This study demonstrates that UnTANGLeD outperforms conventional clustering approaches by integrating multi-resolution consensus building with systematic cluster stratification based on internal coherence, robustness, and biological conservation. Traditional methods often analyse data at a single granularity level, producing a fixed snapshot of biological complexity while suffering from sensitivity to parameter choices, noise-induced artifacts, and inability to determine the optimal number of clusters. Current consensus clustering methods aim to improve sensitivity and robustness through (1) iteratively applying the same or multiple algorithms with varying hyperparameters such as distance metrics^5,8^ (2) clustering repeatedly on subsampled or perturbed data^10,11^ (3) combining results from multiple algorithms^12,13^, or (4) leveraging deep learning based approach such as network fusion^22^. Unfortunately, these methods predominantly optimise at a fixed resolution or discrete granularity level, essentially taking different views of the same biological structure rather than exploring how stable the structures are across a continuous spectrum of granularity. Critically, existing methods lack a systematic framework to evaluate final cluster quality and prioritize promising clusters for follow up, limiting their utility in screening applications where identifying the most promising biological signals is a priority. UnTANGLeD addresses these limitations by leveraging co-occurrence patterns across 100 increasing granularity levels to build a consensus that preserves robust relationships across multiple data scales, while providing metrics to prioritize the most significant clusters, enabling efficient identification of meaningful patterns in high-dimensional screening datasets. To facilitate broad adoption, we provide UnTANGLeD as a user-friendly R package, offering a versatile clustering and prioritisation workflow to guide actionable insights from any highly complex dataset with applications spanning drug discovery, target identification, disease subtyping, biomarker selection with potential for application in non-biological contexts.

## Acknowledgements

This work has been supported by grant funding from the NHMRC (MRFCDDM000033 to NP and 2007625 to NP) and the National Heart Foundation of Australia (106721 to NP). The Victorian Centre for Functional Genomics RRID:SCR_025582 (K.J.S.) is funded by the Australian Cancer Research Foundation (ACRF), Phenomics Australia, through funding from the Australian Government’s National Collaborative Research Infrastructure Strategy (NCRIS) program, the Peter MacCallum Cancer Centre Foundation and the University of Melbourne Collaborative Research Infrastructure Program. KIP acknowledges funding from the Victorian Cancer Agency (Mid-Career Low Survival Cancer Philanthropic Research Fellowship, APP MCRF23014) and the Department of Defense (Ovarian Cancer Research Program Pilot Award, APP OC220022). KJS, IGC, and DC acknowledge support from the Medical Research Future Fund (APP1200264) and Therapeutic Innovation Australia through the Pipeline Accelerator Scheme. DC further acknowledges funding from the Victorian Cancer Agency (MCRF19046) and the Ovarian Cancer Research Foundation (GA-2024-02). M.S.C and D.C would like to acknowledge additional support from the BC Cancer Foundation, Vancouver General Hospital/UBC Hospital Foundation, the Janet D. Cottrelle Foundation, Cure Our Ovarian Cancer, Ovarian Cancer Canada/OVCAN, and the Cancer Research Society. We gratefully acknowledge the patients and their families, including the MacKenzie, Lawler, MacRae, Ho, Luther, Ludemann, and Schmid families. We also thank Professor John Hooper (Mater Research) and the Australian Ovarian Cancer Study for generating and providing cell lines used in this study. We thank Jennii Luu and Robert Vary from the Victorian Centre for Functional Genomics for their screening support. We thank Compounds Australia for the provision of compounds and logistical support. We also thank Sophie Shen for her assistance with the schematic in Figure 1.

## Code Availability

The code used in this study is available upon request.

