## Supplemental Data and Methods for "A scalable, multi-resolution consensus clustering approach for prioritising robust signals from high-throughput screens"

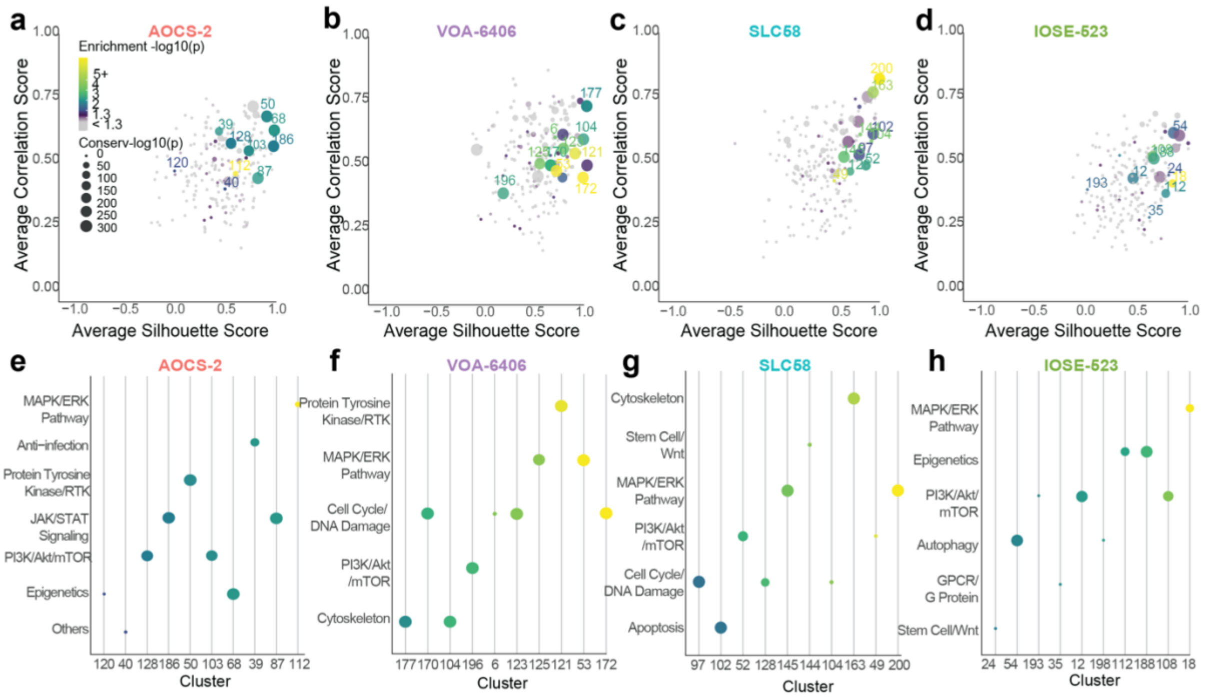


**Extended Figure 1: Unsupervised clustering reveals biologically meaningful drug clusters across ovarian cell lines. (a-d)** Scatter plot of UnTANGLeD clusters in 3 low-grade serous ovarian cancer cell lines (AOCS-2, VOA-6406, SLC58) and 1 normal ovarian surface epithelial cell line (IOSE-523) stratified by average correlation and average silhouette score. (**e-h**) Pathway enrichment for the top 10 clusters with the highest Enrichment -log10 p-adjust in AOCS-2, VOA- 6406, SLC58, and IOSE-523. Enrichment calculated using hypergeometric test. (**a-h**) Size indicates conservation significance (-log10 adjusted p-value) across cell lines while colour represents pathway enrichment significance (-log10 adjusted p-value).

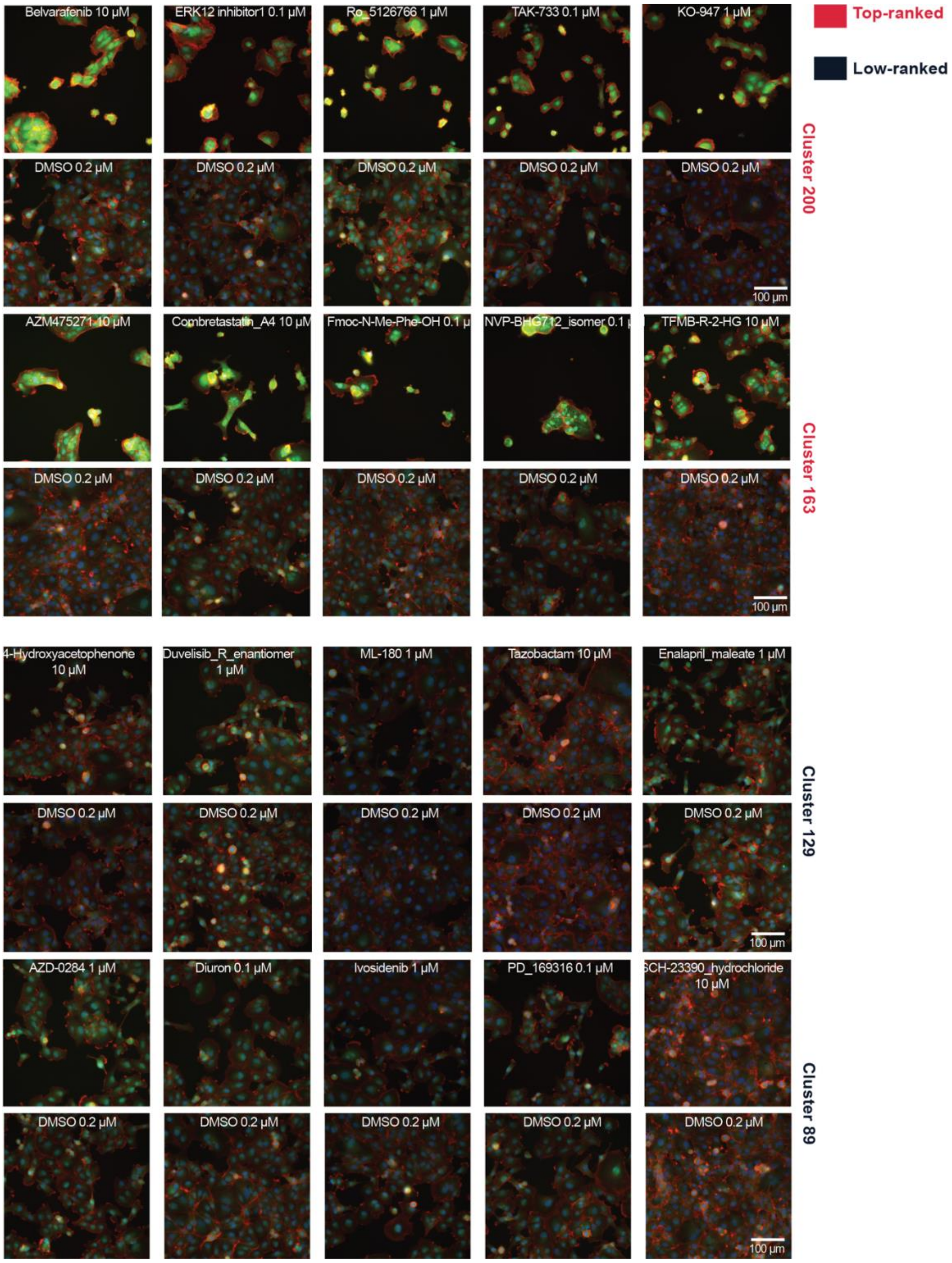


**Extended Figure 2: Representative images of drug-induced morphological changes in SLC58 cells for two example top-ranked and low-ranked UnTANGLeD clusters.** Representative high-content microscopy images of SLC58 low-grade serous ovarian cancer cells treated with various drugs from two top-ranked clusters (200 and 163) and two low-ranked clusters (129 and 89) identified by UnTANGLeD analysis. Each drug treatment is shown alongside its corresponding DMSO control (0.2 µM). Cells were stained with DAPI (blue, nuclei), CellMask (red, plasma membrane), and phalloidin/rhodamine (green, F-actin). Scale bar: 100 µm.

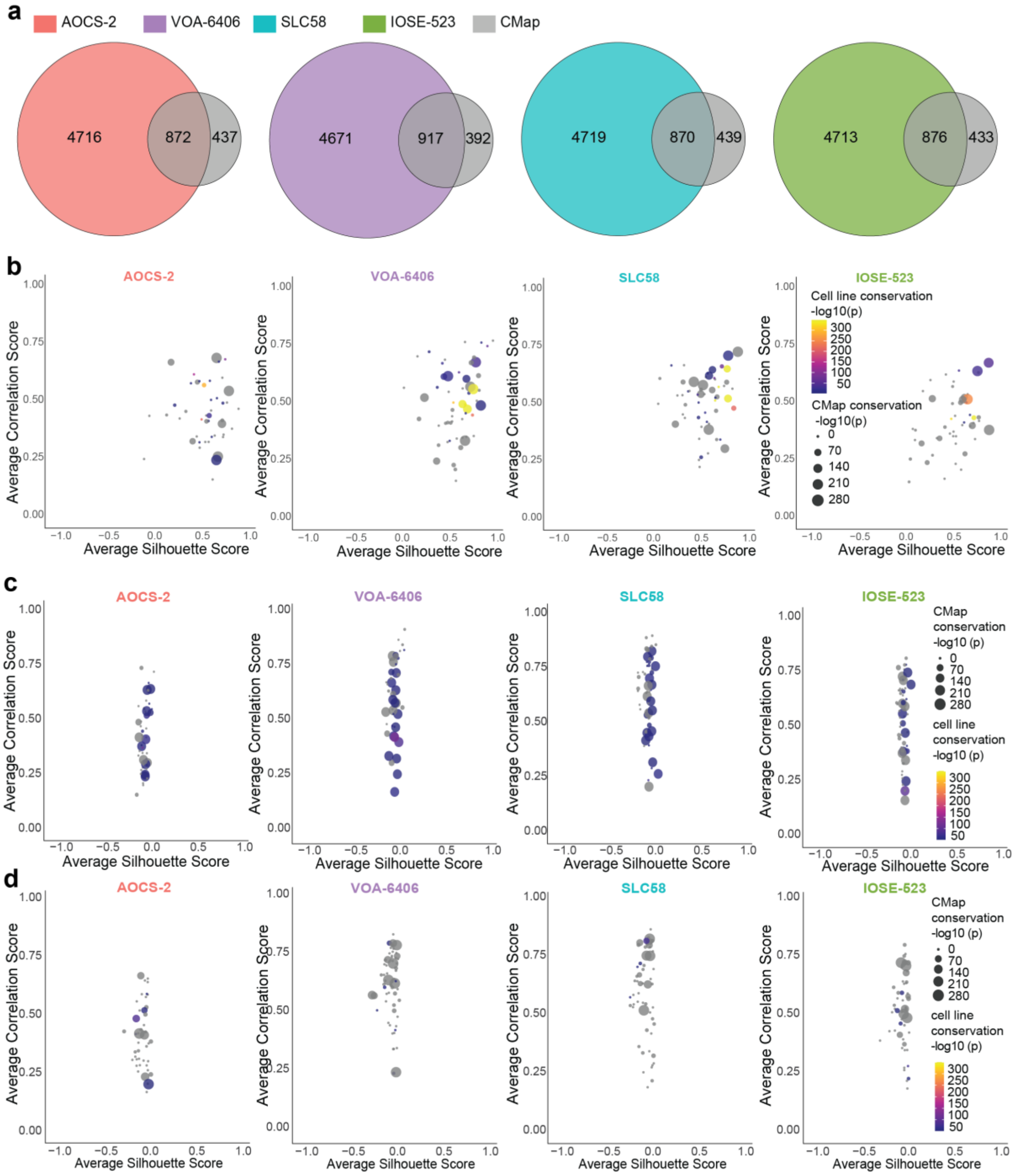


**Extended Figure 3: Orthogonal conservation between morphology-based drug clusters and CMap gene-expression based drug clusters.** (**a**) Overlap between drug compounds after cytotoxicity filtering from 3 low-grade serous ovarian cancer cell lines (AOCS-2, VOA-6406, SLC58) and 1 normal ovarian surface epithelial cell line (IOSE-523) with CMap drug compounds. Conservation of drug clusters from (**b**) UnTANGLeD, (**c**) Hierarchical, and (**d**) K-means clusters for ovarian cell lines AOCS-2, VOA-6406, SLC58, and IOSE-523, stratified by average silhouette score (x-axis) and correlation score (y-axis), limited to clusters containing more than 5 overlapping drug compounds with CMap. (**b-d**) Point size indicating the strength of conservation with CMap clusters (-log10 CMap p.adjust). Colour intensity represents the strength of conservation across ovarian cell lines (-log10 conservation p.adjust).

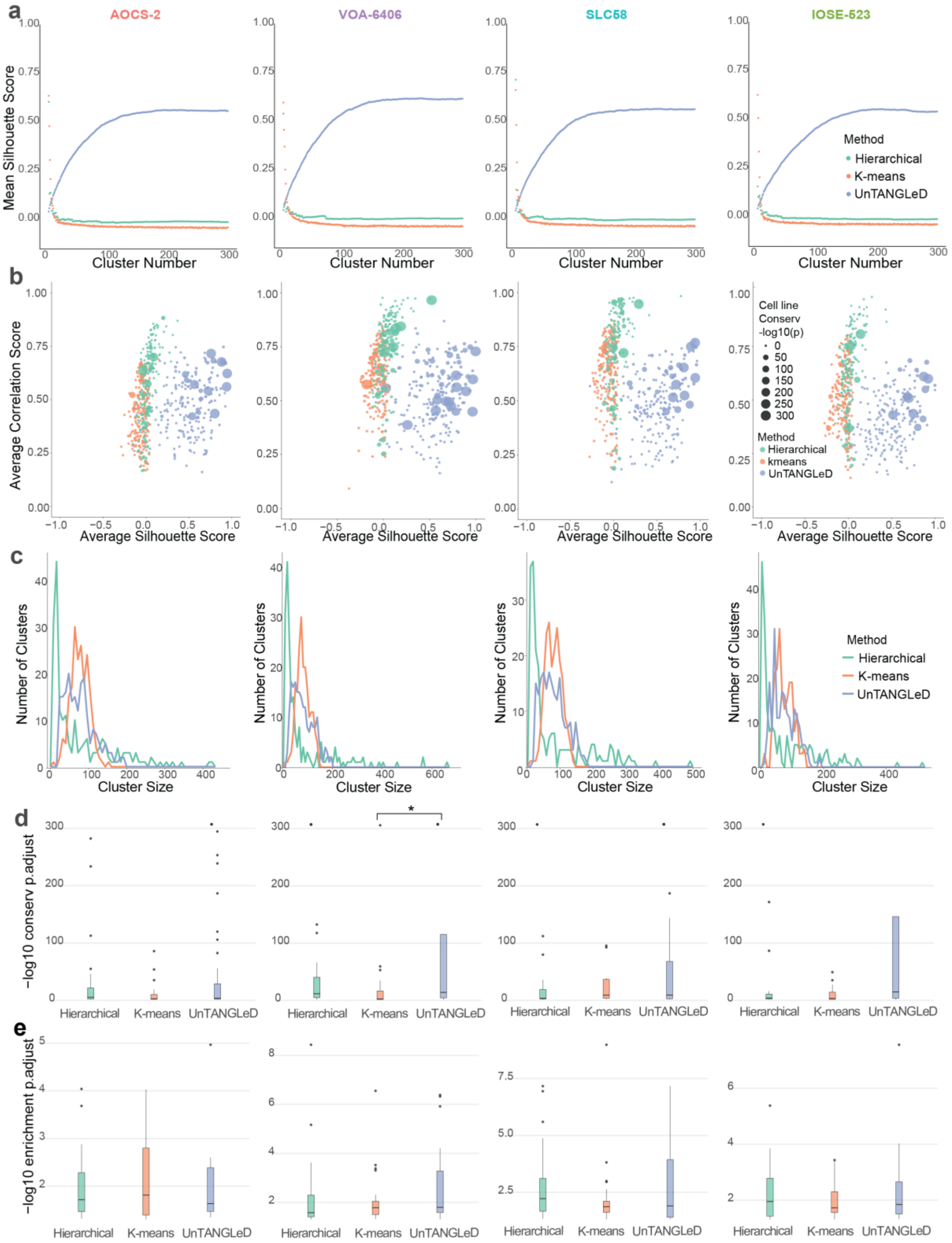


**Extended Figure 4: UnTANGLeD outperforms traditional clustering methods in identifying robust drug clusters. (a)** Comparison of average silhouette scores across cluster numbers (2-300) for UnTANGLeD, hierarchical, and k-means clustering in 3 low-grade serous ovarian cancer cell lines (AOCS-2, VOA-6406, SLC58) and 1 normal ovarian surface epithelial cell line (IOSE-523). Hierarchical and k-means clustered on centred normalised drug-imaging feature matrix and drug-principal component matrix respectively. **(b)** Scatter plot comparing cluster quality metrics (average silhouette score and correlation score) for each method across cell lines. Dot size indicates conservation significance across cell lines (-log10 p- adjust). **(c)** Distribution of cluster sizes for each method across cell lines. **(d)** Violin plots comparing conservation significance (-log10 p-adjust) for clusters with cross-cell line conservation adjusted p-value < 0.05. Statistical significance between pairwise comparison is derived from non-parametric Dunn test. * indicates p < 0.05. **(e)** Violin plots comparing pathway enrichment significance (-log10 p-adjust) for clusters with enrichment adjusted p-value < 0.05. (**a-e**) Colours indicate clustering method (blue: UnTANGLeD, green: hierarchical, orange: k-means). Each column represents results for a different cell line (AOCS-2, VOA-6406, SLC58, IOSE-523), as labelled at the top of the figure.

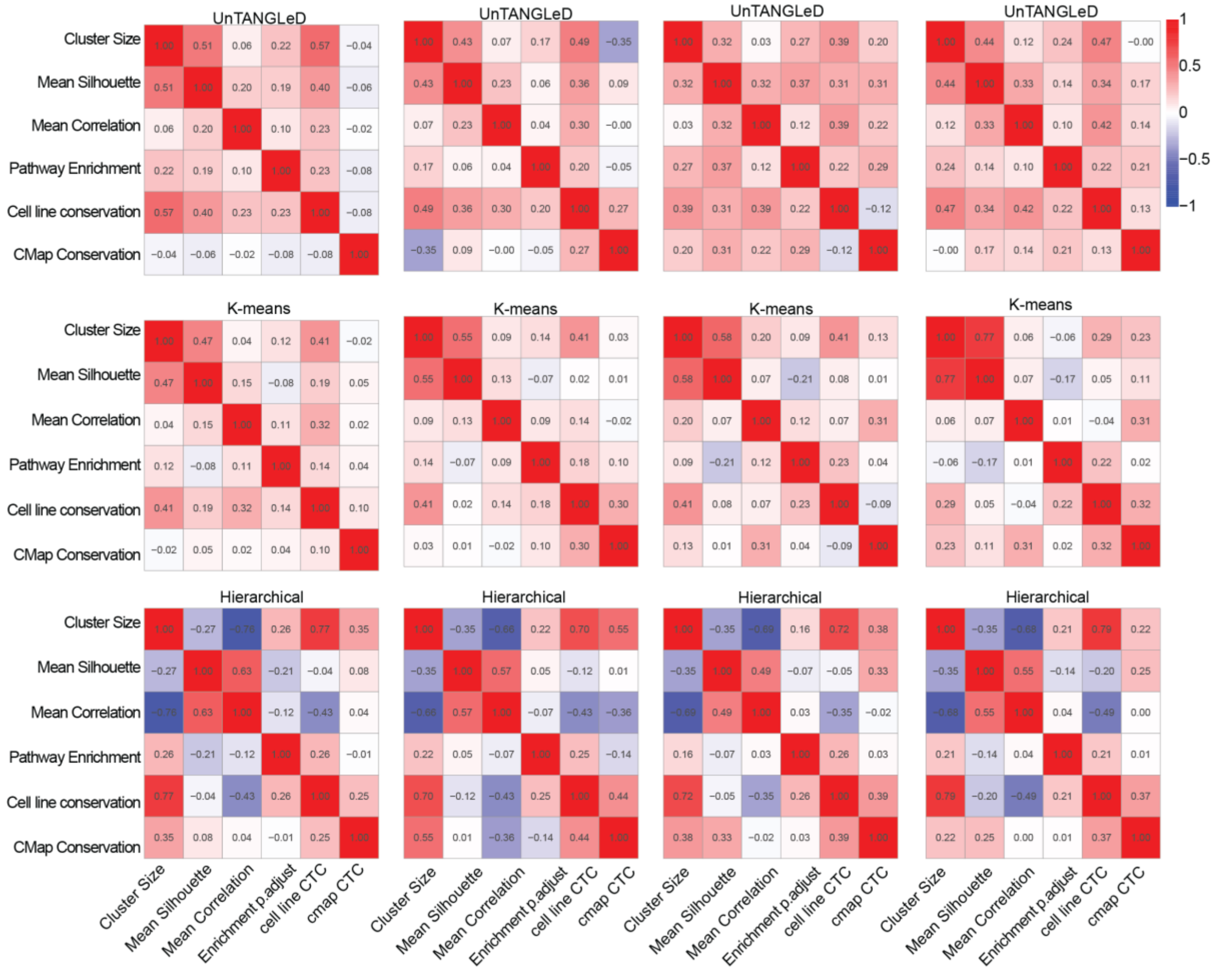


**Extended Figure 5: Spearman correlation of clustering metrics across UnTANGLeD, k-means, and hierarchical clustering methods.** The figure compares the relationships between various clustering metrics for UnTANGLeD, k-means, and hierarchical clustering methods across four ovarian cell lines: AOCS-2, VOA-6406, SLC58, and IOSE-523. Heatmaps showing Spearman correlation coefficients between clustering metrics for each method and cell line. Metrics include cluster size, average silhouette score, average correlation score, pathway enrichment significance (-log10 p-adjust), cell line conservation significance (-log10 p-adjust), and CMap conservation significance (-log10 p-adjust). Colour intensity represents the strength of correlations.

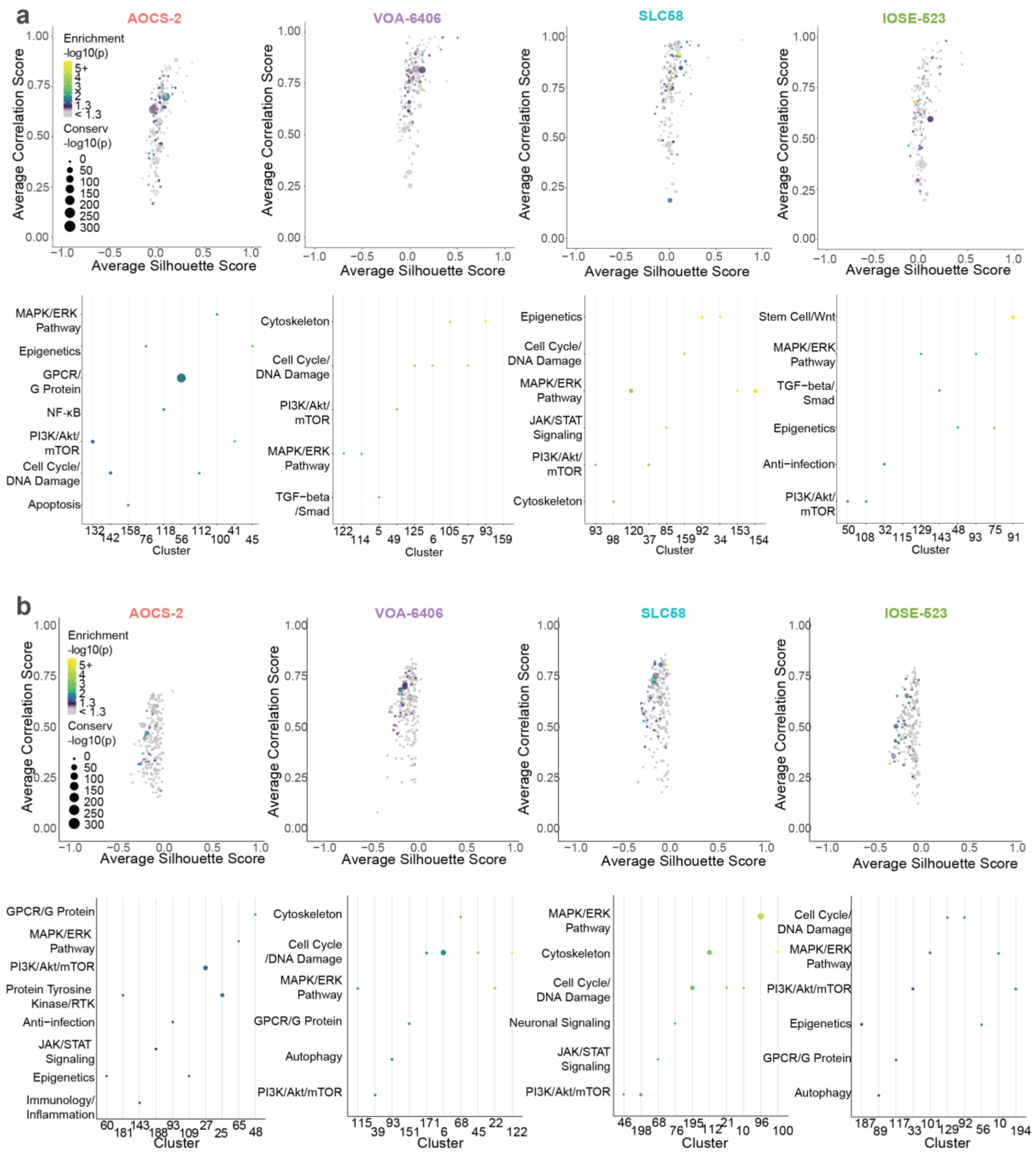


**Extended Figure 6**: **Hierarchical, and k-means cluster silhouette-correlation plots with enriched pathway in top 10 clusters.** Clusters from (**a**) hierarchical and (**b**) k-means across four ovarian cell lines: AOCS-2, VOA-6406, SLC58, and IOSE-523. stratified by average silhouette score (x-axis) and average correlation score (y-axis). Pathway enrichment for the top 10 clusters with the highest enrichment significance (-log10 p-adjust) for each cell line. (**a-b**) Point size indicates pathway enrichment significance (-log10 p-adjust) and colour intensity indicates conservation significance across cell lines (-log10 p-adjust).

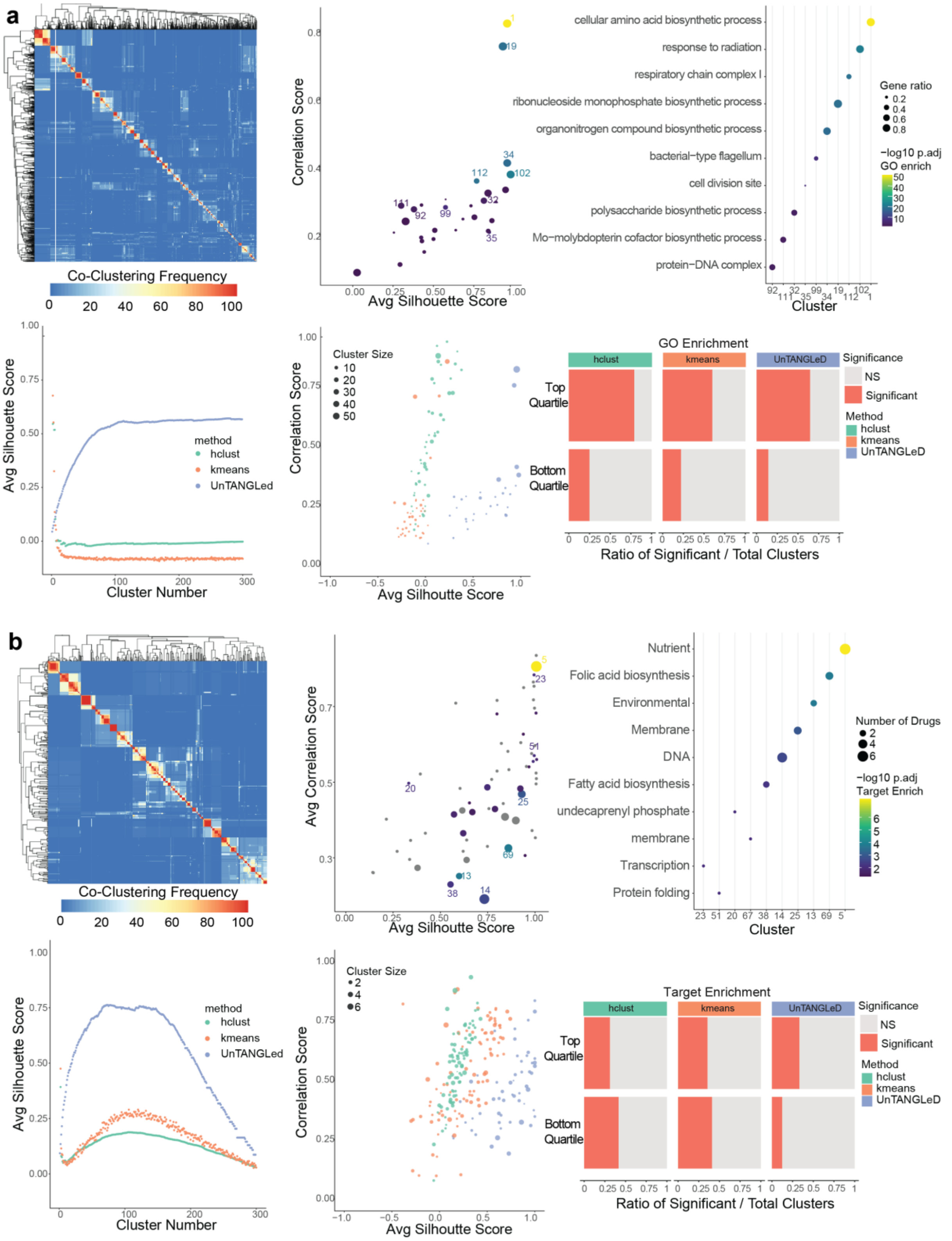


**Extended Figure 7**: **Unsupervised clustering of genes and stress conditions in E. coli single deletion library**

(**a**) Analysis of gene and (**b**) condition clustering results. Consensus matrix heatmap showing co-clustering frequency across resolutions for gene pairs; Scatter plot of UnTANGLeD clusters stratified by average silhouette score and correlation score, with top enriched GO terms annotated. Point size indicates gene ratio and colour represents enrichment significance (-log10 p-adjust); Comparison of average silhouette scores across increasing cluster numbers (2-300) for UnTANGLeD, hierarchical clustering (hclust), and k-means methods; Comparative scatter plots showing cluster distribution for the three clustering methods; Comparison of significant cluster ratios across clustering methods based on three biological relevance metrics. Top panel shows top-performing clusters (>75% quantile) while bottom panel shows bottom-performing clusters (<25% quantile), as defined by 75th and 25th percentile of average correlation and silhouette scores. Biological significance is defined as followed: GO enrichment (adj. p-value <0.05) and target enrichment (adj. p-value <0.05).

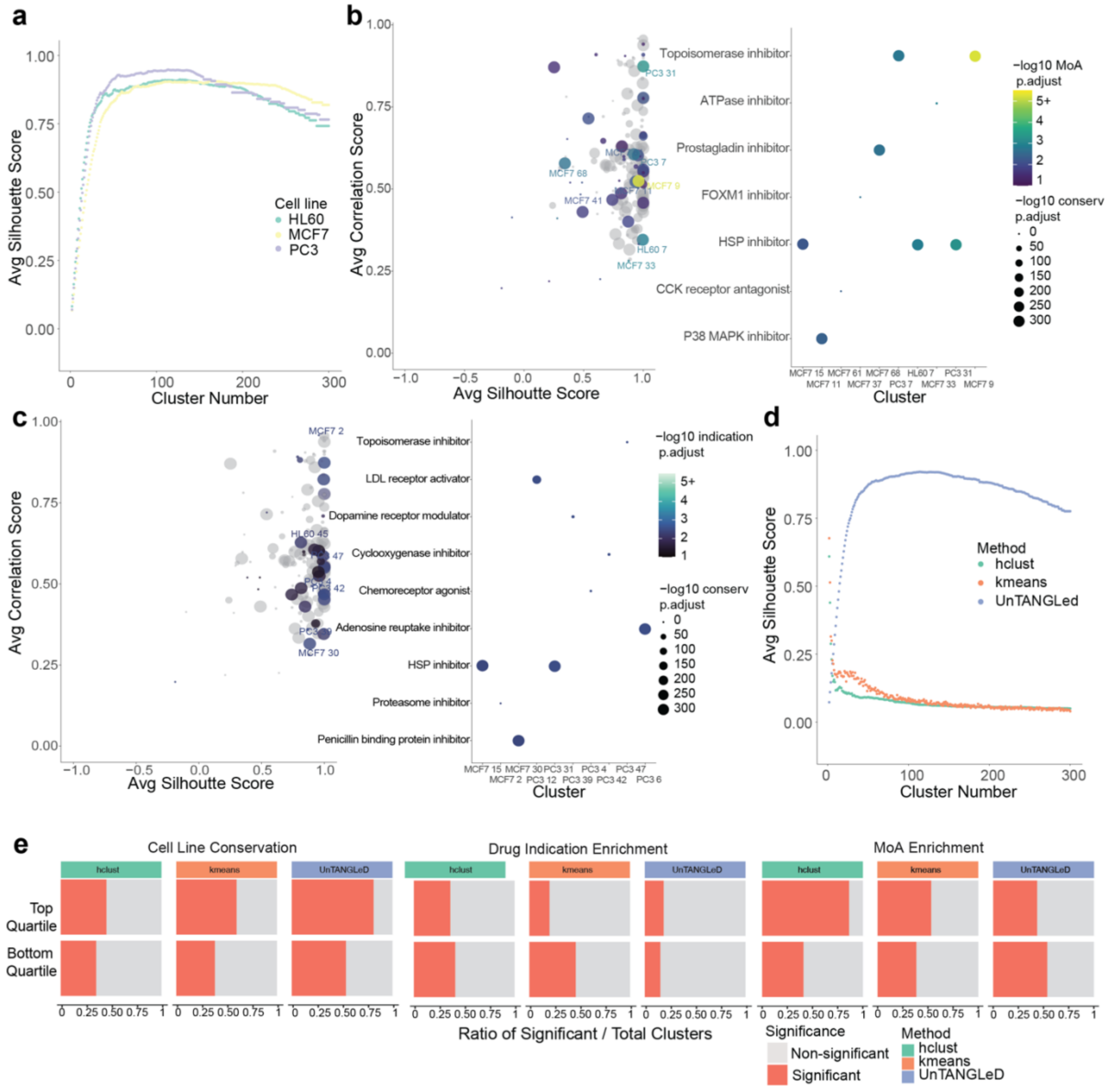


**Extended Figure 8**: **Unsupervised clustering of conditions in Connectivity Map**

(**a**) Average silhouette scores across increasing cluster numbers for three CMap cell lines (HL60, MCF7, and PC3). (**b-c**) Clusters from all CMap cell lines (HL60, MCF7, and PC3), stratified by average silhouette score (x-axis) and correlation score (y-axis). Point size indicates conservation significance across cell lines (-log10 p-adjust). Colour indicates (**b**) mechanism of action (MoA) and (**c**) drug indication enrichment significance (-log10 p-adjust). Moa and drug indications are annotated for the top 10 clusters with the highest enrichment -log10 p-adjust. (**d**) Comparison of average silhouette scores for three CMap cell lines across increasing cluster numbers (2-300) for UnTANGLeD, hierarchical clustering (hclust), and k-means methods (**e**) Comparison of significant cluster ratios across clustering methods based on three biological relevance metrics. Top panel shows top-performing clusters (>75%

quantile) while bottom panel shows bottom-performing clusters (<25% quantile), as defined by 75th and 25th percentile of average correlation and silhouette scores. Significance is defined as followed: Cell line conservation (adj. p-value <0.05), drug indication enrichment (adj. p-value <0.05), and MoA enrichment (adj. p-value <0.05). (**b-e**) Data shown represents the combined or average of HL60, MCF7, and PC3 cell lines.

**Extended Table 1: Pairwise Fisher's Exact Test on Biologically Significant Cluster Ratios Across Clustering Method Pairs**

**a.** Top Quartile Clusters (>75% quantile for silhouette and correlation scores)

| Metric | Method Comparison | Odds Ratio | 95% Confidence Interval | p-value | Adjusted p-value | Significance |
|---|---|---|---|---|---|---|
| Pathway Enrichment | hclust vs. kmeans | 3.54 | 1.12 - 14.87 | 2.42e-02 | 3.78e-02 | * |
| Pathway Enrichment | hclust vs. UnTANGLeD | 0.99 | 0.45 - 2.20 | 1.00e+00 | 1.00e+00 | ns |
| Pathway Enrichment | kmeans vs. UnTANGLeD | 0.28 | 0.06 - 0.93 | 2.84e-02 | 3.78e-02 | * |
| Conservation | hclust vs. kmeans | 2.26 | 0.57 - 13.13 | 2.59e-01 | 3.11e-01 | ns |
| Conservation | hclust vs. UnTANGLeD | 0.10 | 0.04 - 0.21 | 2.71e-11 | 1.70e-10 | **** |
| Conservation | kmeans vs. UnTANGLeD | 0.04 | 0.01 - 0.15 | 2.83e-11 | 1.70e-10 | **** |
| CMap zscore | hclust vs. kmeans | 0.07 | 0.01 - 0.35 | 5.38e-05 | 1.29e-04 | *** |
| CMap zscore | hclust vs. UnTANGLeD | 0.05 | 0.01 - 0.22 | 1.78e-07 | 7.14e-07 | **** |
| CMap zscore | kmeans vs. UnTANGLeD | 0.71 | 0.30 - 1.63 | 4.38e-01 | 4.78e-01 | ns |

**b.** Bottom Quartile Clusters (<25% quantile for silhouette and correlation scores)

| Metric | Method Comparison | Odds Ratio | 95% Confidence Interval | p-value | Adjusted p-value | Significance |
|---|---|---|---|---|---|---|
| Pathway Enrichment | hclust vs. kmeans | 0.86 | 0.34 - 2.27 | 8.25e-01 | 9.90e-01 | ns |
| Pathway Enrichment | hclust vs. UnTANGLeD | 1.88 | 0.64 - 6.30 | 2.41e-01 | 3.61e-01 | ns |
| Pathway Enrichment | kmeans vs. UnTANGLeD | 2.18 | 0.68 - 7.76 | 1.91e-01 | 3.43e-01 | ns |
| Conservation | hclust vs. kmeans | 4.31 | 0.90 - 41.17 | 4.83e-02 | 1.45e-01 | ns |
| Conservation | hclust vs. UnTANGLeD | 3.03 | 0.77 - 17.51 | 1.01e-01 | 2.42e-01 | ns |
| Conservation | kmeans vs. UnTANGLeD | 0.71 | 0.06 - 6.40 | 1.00e+00 | 1.00e+00 | ns |
| CMAP zscore | hclust vs. kmeans | 3.04 | 1.02 - 11.06 | 4.16e-02 | 1.45e-01 | ns |
| CMAP zscore | hclust vs. UnTANGLeD | 1.94 | 0.75 - 5.51 | 2.00e-01 | 3.43e-01 | ns |

| | | | | | | |
|---|---|---|---|---|---|---|
| CMAP zscore | kmeans vs. UnTANGLeD | 0.64 | 0.15 - 2.39 | 5.61e-01 | 7.49e-01 | ns |

**** p < 0.0001, *** p < 0.001, ** p < 0.01, * p < 0.05, ns = not significant; Combined results from all four cell lines (AOCS-2, VOA-6406, SLC58, IOSE-523); Biological significance of clusters defined as Pathway Enrichment (adj. p-value < 0.05), Cross-cell line conservation (adj. p-value < 0.05), CMap z-score (|z| > 2), CMap p-adjust (adj. p-value < 0.05)

**Extended Table 2: Pairwise Fisher's Exact Test on Biologically Significant Cluster Ratios Between Top and Bottom Quartiles**

| Metric | Method | Odds Ratio | 95% Confidence Interval | p-value | Adjusted p-value | Significance | Top Significant | Top Not Significant | Bottom Significant | Bottom Not Significant |
|---|---|---|---|---|---|---|---|---|---|---|
| Pathway Enrichment | UnTANGLeD | 2.15 | 0.74 - 7.16 | 1.62e-01 | 2.42e-01 | ns | 16 | 68 | 6 | 55 |
| Conservation | UnTANGLeD | 21.86 | 6.33 - 117.99 | 8.87e-11 | 1.06e-09 | **** | 45 | 39 | 3 | 58 |
| CMAP zscore | UnTANGLeD | 2.34 | 0.91 - 6.60 | 6.36e-02 | 1.27e-01 | ns | 22 | 62 | 8 | 53 |
| Pathway Enrichment | kmeans | 0.28 | 0.06 - 1.01 | 5.02e-02 | 1.21e-01 | ns | 4 | 61 | 11 | 46 |
| Conservation | kmeans | 1.33 | 0.15 - 16.43 | 1.00e+00 | 1.00e+00 | ns | 3 | 62 | 2 | 55 |
| CMAP zscore | kmeans | 2.58 | 0.79 - 9.93 | 1.24e-01 | 2.12e-01 | ns | 13 | 52 | 5 | 52 |
| Pathway Enrichment | hclust | 1.13 | 0.52 - 2.55 | 8.53e-01 | 9.31e-01 | ns | 21 | 90 | 15 | 73 |
| Conservation | hclust | 0.70 | 0.26 - 1.83 | 5.05e-01 | 6.05e-01 | ns | 11 | 100 | 12 | 76 |
| CMAP zscore | hclust | 0.06 | 0.01 - 0.27 | 2.94e-06 | 1.76e-05 | **** | 2 | 109 | 20 | 68 |

**** p < 0.0001, *** p < 0.001, ** p < 0.01, * p < 0.05, ns = not significant; Combined results from all four cell lines (AOCS-2, VOA-6406, SLC58, IOSE-523); Biological significance of clusters defined as Pathway Enrichment (adj. p-value < 0.05), Cross-cell line conservation (adj. p-value < 0.05), CMap z-score (|z| > 2), CMap p-adjust (adj. p-value < 0.05)
